# Contrasting effects of aging on the expression of transposons, the piRNA machinery and mitochondrial transcripts in the *Drosophila* ovary

**DOI:** 10.1101/342105

**Authors:** Alexandra A. Erwin, Justin P. Blumenstiel

## Abstract

Redistribution of heterochromatin during aging has been linked to the de-repression of transposable elements and an overall loss of gene regulation in the soma. Whether or not epigenetic factors such as heterochromatin marks are perturbed in reproductive and germline tissues is of particular interest because some epigenetic factors are known to transmit across generations. Additionally, the relative contribution of factors intrinsic or extrinsic to the germ line have in reproductive decline remains unknown. Using mRNA sequencing data from late stage egg chambers in *Drosophila melanogaster*, we show that age-related expression changes occur in genes residing in heterochromatin, particularly on the largely heterochromatic 4^th^ chromosome. In addition, we identify an increase in expression of the piRNA machinery. We further identify a striking age-related reduction in mitochondrial transcripts that we can attribute to the somatic tissues. Other than a modest increase in overall TE expression in the aging germline, we find no global TE de-repression in reproductive tissues. Rather, the observed effects of aging on TEs are primarily strain and family specific. These results indicate unique responses in somatic versus germline tissue with regards to epigenetic aging effects and suggest that the global loss of TE control observed in other studies may be specific to certain tissues, genetic backgrounds and TE family. This study also demonstrates that while age-related effects can be maternally transmitted, the germline is generally robust to age-related changes.

## INTRODUCTION

The age-related decline of the reproductive system has important consequences for evolution because reproductive success determines the fitness of an organism. Since the majority of aging studies focus on overall somatic decline, relatively little is known about the causes of reproductive aging. In humans, progressive delays in childbearing are leading more people to confront the reduced fertility and fecundity that accompanies advanced age (Billari et al., 2007; Dunson et al., 2002). Reproductive senescence is not unique to mammals, however. The invertebrate model *Drosophila melanogaster* shows a progressive decline in egg production at middle age, thought to be partially caused by a reduction in germline stem cell proliferation and decreased survival of developing eggs (Zhao et al., 2008). Possible mechanisms underlying these changes include reduced ovariole number, decreased rates in germline stem cell division, and apoptosis in egg chambers of older females (Pan et al., 2007; Zhao et al., 2008). Animals may have conserved mechanisms to regulate reproductive decline and control the relationship between reproduction and lifespan. Not only have mechanisms of gametogenesis been found to be similar across organisms, but the control of ovulation has also been shown to be conserved between *Drosophila* and humans (Sun and Spradling, 2013). Because *Drosophila* is an established model for studies of both reproductive and somatic aging, we used it here to examine age-related genome-wide expression changes in the germline and broader reproductive tissues.

While genetic causes have long been shown to determine longevity - through either inherited or somatic mutation, non-genetic contributions are also proving to be major factors. Epigenetic chromatin marks play an essential role in the maintenance of genome integrity through their repression of genes, repeat sequences, and transposable elements (reviewed by (Putiri and Robertson, 2011). The misregulation of epigenetic marks has been associated with many diseases, including kidney disease, nuerodegenerative diseases, and cancer (Figueroa-Romero et al., 2012; Muntean and Hess, 2009; Smyth et al., 2014). Recently, epigenetic mis-regulation has been attributed to playing a key role in the aging process. In particular, the landscape of silent heterochromatin has been shown to redistribute in aged stem cells and cells of the soma, leading to aberrant gene expression (Bell et al., 2012; De Cecco, 2013a; Jiang, 2013; Larson et al., 2012; Shah et al., 2013; Wood et al., 2010). An additional consequence of this redistribution of heterochromatin is the observed de-repression of transposable elements in the soma during aging, notably in brains and fat body of *Drosophila*, and in a variety of other organisms including mammals (Chen et al., 2016; De Cecco, 2013b; Li et al., 2013; Maxwell et al., 2011; Patterson et al., 2015). Although interesting for the biology of aging, somatic cells do not affect future generations. Surprisingly, little is known about whether epigenetic changes occur in aged reproductive tissues and germline cells that may transmit these non-genetic but potentially heritable effects to the next generation.

The germ line is considered an immortal cell lineage. Thus, germ cells have unique strategies to faithfully transmit DNA indefinitely, such as greater telomerase maintenance (Wright et al., 1996) and greater resistance to genotoxic stress than somatic cells (Vinoth et al., 2008). However, age-related changes in the germline are known to occur. For example, some germ cells lose the ability to divide and differentiate normally (Zhao et al., 2008), the sperm of older human males are thought to be at risk for more de novo mutations based on parent-offspring mutations (Kong et al., 2012), and double strand break repair in oocytes in humans and mice declines with age (Titus et al., 2013). Additionally, age-dependent meiotic nondisjunction may be due to a loss of the protein complex that regulates the separation of sister chromatids over time (Subramanian and Bickel, 2008). However, some age-effects that have been observed in the germline may be due to extrinsic factors such as the microenvironment of the germ line stem cells (Boyle et al., 2007; Pan et al., 2007; Zhao et al., 2008). The relative roles of extrinsic versus intrinsic factors in contributing to germline aging are still being explored. In mammals, much of the current evidence points to a greater role of cell-extrinsic factors. Similar to flies, niche deterioration also may play a role in the mammalian system (Zhang et al., 2006). For example, it has been shown that mammalian spermatagonial stem cells, when transplanted to a young environment, have extended functionality (Ryu et al., 2006; Schmidt et al., 2011). Signaling factors like insulin may also play a role in maintaining germline function in mammals (Hsu and Drummond-Barbosa, 2008; Yang et al., 2013). Thus, while the germline is generally considered to be immortal, components of the germline and its microenvironment are not immune to age-related changes.

Recent findings highlighting the large role of epigenetic changes in the aging process leads us to question whether similar mechanisms may also be at play in reproductive tissues. Although the majority of epigenetic marks are erased and re-established between generations, some epigenetic modifications are transmitted across generations through the germline. Longevity itself is a trait that has been shown to be epigenetically inherited in *C. elegans* (Greer et al., 2016; Greer et al., 2011; Spracklin et al., 2017). Of most relevance, *Drosophila* oocytes transmit the repressive histone mark H3K27me3 to their offspring (Zenk et al., 2017). This creates a potential for age-effects to be passed on to the next generation, an outcome that could pose new questions for traditional evolutionary aging theories that have been around for decades.

Few studies have characterized genome-wide, age-related expression in ovaries and we are not aware of any such studies in the *Drosophila* germline. We sought to determine whether age-related epigenetic changes occur in the germline and broader *Drosophila* reproductive tissues using mRNA expression as a proxy. Specifically, we asked whether the age-dependent transposable element release extends to the ovary by determining whether transposable elements were derepressed during aging. We further tested the heterochromatin aging hypothesis by testing whether genes in or near heterochromatin boundaries were aberrantly expressed, and if genes were globally misregulated in reproductive tissues. We find that gene expression changes are enriched in heterochromatic regions of the genome, but the direction of change is not consistent with a global increase in expression of heterochromatin. Further, we only find idiosyncratic aging effects on TE expression and no global increase in expression. These results suggest that the age-related transposon release and the heterochromatin aging hypothesis do not extend to the *Drosophila* ovary in a simple manner.

## METHODS

### Fly stocks

*D. melanogaster* DGRP lines 237 and 321 were utilized for this study and maintained at 22 degrees Celsius and 12 hour light cycles.

### Egg Chamber Tissue Collection

Flies were maintained in bottles at controlled larval density (∼100 per bottle) for two generations before tissue collections. Zero to one day old F2 females were transferred to individual vials for aging treatment and supplemented with two males ranging from 3-7 days old approximately every seven days to encourage egg production. Flies were moved to fresh vials weekly. Stage 14 egg chambers were dissected from ovaries of 3-4 and 32-34 day old females in PBS buffer. Using a thinly bristled paintbrush, 2-5 egg chambers from each female were added to single caps of .2mL tubes, stabbed with RNAse free needles in 30uL TRIzol, and flash frozen in liquid nitrogen.

### Embryo Tissue Collection

Embryos were collected from the Ral-321 strain only. Flies were maintained in bottles at controlled larval density (∼100 per bottle) for two generations. F2 females were maintained continuously laying in bottles containing yeast paste and supplemented with younger males. For embryo collections, flies were moved to mating cages with petri dishes filled with fly food and ∼5mL of yeast paste to acclimate overnight. Embryos were plucked from food plates after approximately 45 minutes of laying. Embryos were rinsed with embryo wash (0.7%NaCl, 0.05% Triton X-100) and dipped into 50% bleach using a mesh net for 30s-1min followed by another rinse with embryo wash. Embryos were picked up with a thinly bristled brush and put into a TRIzol filled .2mL tube cap and flash frozen in liquid nitrogen.

### RNA extraction and mRNA Sequencing

For RNA extraction, egg chambers from 5 females (∼20 egg chambers total) were pooled. In total there were 5 pools for each age treatment across both strains. For embryos, ∼20 embryos from each cage were pooled with 4 cages across two timepoints. Accounting for the TRIzol already in the samples from the collection stage, we added up to a total volume of 300uL TRIzol for RNA extractions. To improve recovery in the separation phase, we used 5PRIME Phase Lock Gel Heavy tubes. RNA was resuspended in 25uL of H20. Library preps were performed using the NEBNext Ultra Kit according to the manufacturer’s instructions (New England Biolabs). NEBNext Ultra libraries were pooled in groups of 8-10 per lane, and run with single-end 100 bp reads on a HiSeq 2500.

### Analysis of mRNA sequencing data

RNA-seq was performed in CLC Genomics Workbench 8 using release 6 of the *Drosophila melanogaster* reference genome. For expression values, RPKM estimates generated by the RNA-seq tool in CLC Genomics Workbench were used. FDR-adjusted p-values for significant differential expression were calculated with a CLC algorithm based on the DESeq2 package in Bioconductor (Love et al., 2014). To estimate TE family expression, an annotated TE library was included in the RNAseq analysis while the rest of the genome was masked for individual TE sequences. GO analysis was performed with GOrilla (Eden et al., 2009) using *D*. *melanogaster* orthologs genes sorted by FDR p-value for the test of treatment effect.

## RESULTS

### Genic transcripts differentially expressed with age in egg chambers of both strains

A number of studies have compared aging transcriptomes across tissues and even across species (Doroszuk, 2012; Lee, 1999; McCarroll et al., 2004; Pletcher, 2002; Zhan et al., 2007; Zou et al., 2000). Fewer studies, however, compare profiles in more than one natural strain (Highfill et al., 2016; Landis et al., 2004). Here we sought to determine how gene expression is modulated in the aging ovary in two different inbred Raleigh strains of *Drosophila melanogaster* obtained from the DGRP (Mackay et al., 2012). Since ovaries are highly heterogeneous, consisting of a mixture of somatic tissues, germline-stem cells and many different stages of oogenesis, we focused our RNAseq analysis using stage 14 egg chambers. This allowed us to minimize variation of cell type composition and to enrich for age-effects in the germline. Stage 14 egg chambers consist of an oocyte surrounded by a follicular sheath and represent the last stage of oogenesis before fertilization and oviposition. To measure differences in gene expression, we compared expression profiles in stage 14 egg chambers from mothers at 3-4 and 32-34 days post-eclosion (sample overview presented in Table 1). Overall, we identified 300 transcripts that were differentially expressed (DE) between young and old stage-14 egg chamber samples in a combined analysis with the two Raleigh lines (FDR adjusted p-value <.05), testing for age while controlling for strain in DESeq2.

**Table 3. 1.** Sample overview of stage 14 egg chambers. Ral 237 and Ral 321 were the two DGRP strains utilized for RNA sequencing analysis. D=days post-eclosion. Each biological replicate is a pool of egg chambers from five females.

| DGRP STRAINS | # BIOLOGICAL<br>REPLICATES (3-4D) | # BIOLOGICAL<br>REPLICATES (32-34D) |
| --- | --- | --- |
| RAL_321 | 4 | 4 |
| RAL_237 | 3 | 3 |

Of the DE transcripts identified in the combined analysis, 106 transcripts show an average increase with age, while 194 show an average decrease with age across strains (Fig. 3.1A). Figure 3.1B demonstrates that the significantly differentially expressed transcripts are strongly correlated and show the same direction of expression changes between old and young egg chambers across the two strains (Pearson’s product-moment correlation = 0.66, p-value < 2e-16). Seven of these genes have previously been associated with regulation of lifespan. Notably, *hebe* (CG1623) overexpression increases both longevity and fecundity (Li and Tower, 2009) and *Hsp27* overexpression increases lifespan (Wang et al., 2004). Both of these transcripts showed average lower expression in older stage-14 egg chambers across the two strains (*hebe*: 4.11-fold decrease, FDR p-value < .1.28E-05; *Hsp27*: 1.6-fold decrease; FDR p-value <.006). *Hsp27* was also one of the most highly expressed genes (26th). Another gene, *POSH* (Plenty of SH3s, CG4909) has been shown to promote cell survival in both *Drosophila* and human cells when overexpressed (Tsuda et al., 2010). We find that this transcript shows a 1.46-fold increase with age in egg chambers (FDR adjusted p-value<4.05E-05). The other transcripts previously associated with regulation of lifespan include Thiolase (CG4581), Thor (CG8846), Coq2 (Coenzyme Q biosynthesis protein 2; CG9613), and Tpi (triose phosphate isomerase; CG2171). Other notable categories of gene ontology analysis using GOrilla (Eden et al., 2009) results for biological process by rank significance include terms pertaining to the electron transport chain (GO:0022900; GO:0022904), mitochondrial electron transport chain (GO:0006120), numerous metabolic processes, developmental and cellular processes involved in reproduction (GO:0003006; GO:0022412), eggshell chorion assembly (GO:0007306), many terms related to regulation of mitochondrial organization and fusion, determinant of adult life span (GO:0008340) and interestingly, miRNA metabolic process (GO:0010586). Full results from a gene ontology (GO) analysis for biological process, component, and function by rank significance is shown in Supplementary Table 3.1.

**Figure 3. 1.**
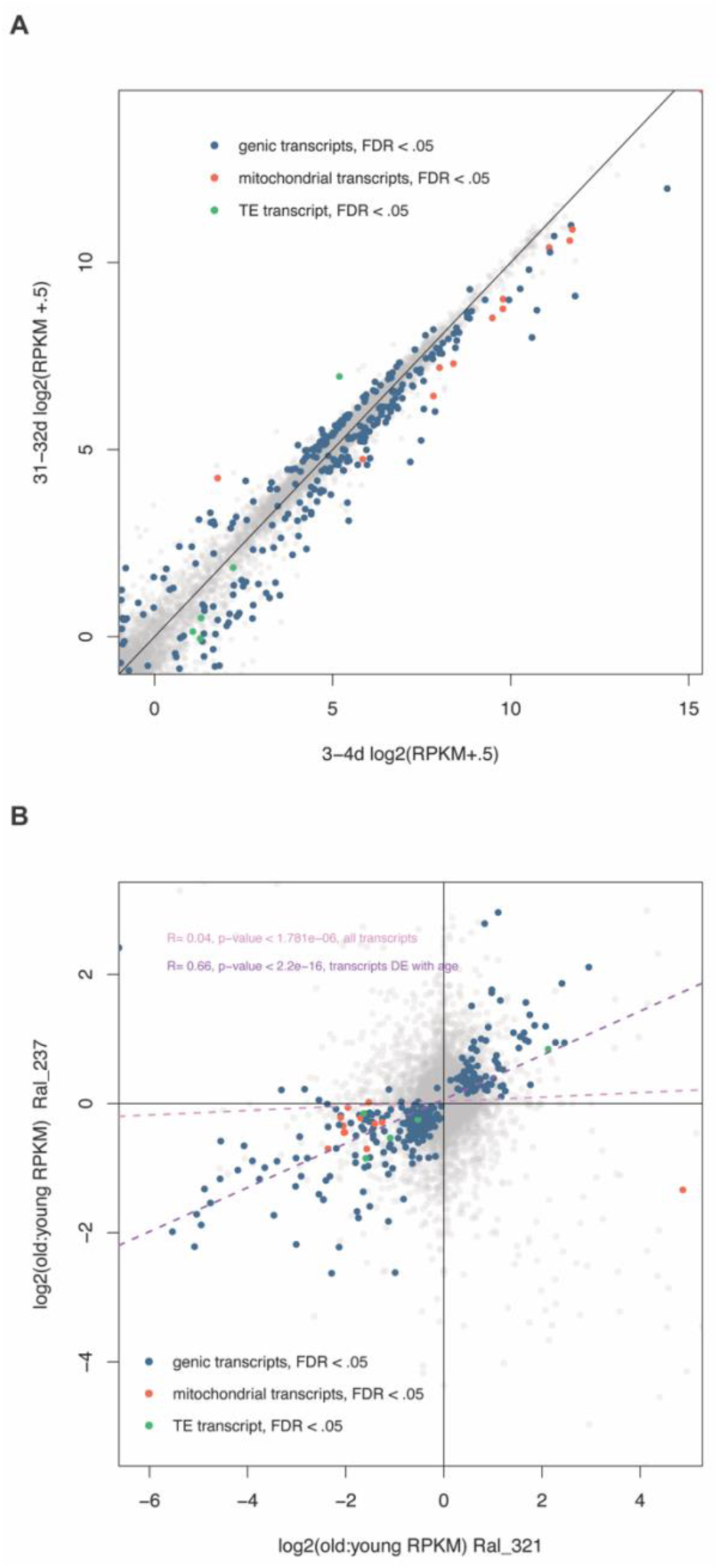
Signature of age-related expression in egg chambers across genetic background. (A) Average log 2(RPKM+.5) expression of stage 14 egg chamber transcripts of old 30 - 34 day old samples versus young 3-4 day old samples. Transcripts significantly expressed between young and old in a paired analysis (FDR<.05) are colored according to transcript type. Five TE transcripts are significantly differentially expressed across both strains with age, with only one, copia, showing an increase in expression. (B) Log2 ratios of old to young (RPKM+.5) expression between strains. The differentially expressed transcripts (FDR p <.05) are strongly and significantly correlated with age across strains.

While these DE transcripts may provide a signature of senescence for egg chambers, the transcriptome, as a whole, shows only a very weak correlation in age-related patterns of expression across these two strains (Pearson’s product-moment correlation = 0.04, p-value < 1.8e-06, Figure 3.1B). This demonstrates that many observed changes in gene expression in the aging ovary are likely to be strain specific. In fact, 59 genes show a significant strain by age effect in our analysis.

### Egg chamber transcripts from the mitochondrial genome are significantly downregulated with age across both strains

Some sets of genes and gene pathways show consistent and concerted changes with age across various studies. Age-related changes in the expression of mitochondrial genes and genes associated with the electron transport chain have consistently been reported. This is most commonly observed as a decrease during aging (Andreu, 1998; Calleja, 1993; Fernandez-Silva SP, 1991; Girardot et al., 2006; Morel, 1995; Sohal et al., 2008). In particular, this pattern has been observed in transcripts associated with the mitochondrial electron transport chain in the gonads of mice (Sharov et al., 2008).

In stage 14 egg chambers, 11 transcripts from the mitochondrial genome significantly decreased with age in the DE analysis (Figure 3.1, Figure 3.2A). Nine of those transcripts also showed a significant strain by age effect, with greater age-related fold-changes observed in Ral_321 for seven transcripts, while two showed opposite age-related effects across the strains (Fig 3.2A). In addition to transcripts from the mitochondrial genome, we also found nuclear transcripts associated with the electron transport chain significantly enriched in a gene ontology analysis (Supp. Table 3.1). All of these nuclear transcripts were also downregulated with age in both strains (Fig. 3.2B). The downregulation of mitochondrial transcripts and those associated with the electron transport chain is in line with established mitochondrial dysfunction associated with age. Our finding lends support to decreased expression of mitochondrial transcripts being a general feature of aging across all tissue types but also highlights strain-specific discrepancies in the magnitude of mitochondrial age-related effects. The reduced expression of mitochondrial transcripts in reproductive tissues may be especially significant as this could contribute to the reduced oocyte quality seen in aged flies (Calleja, 1993; Girardot et al., 2006; Morel, 1995; Sohal et al., 2008) and humans (Johnson et al., 2007; Zhang et al., 2017).

**Figure 3. 2.**
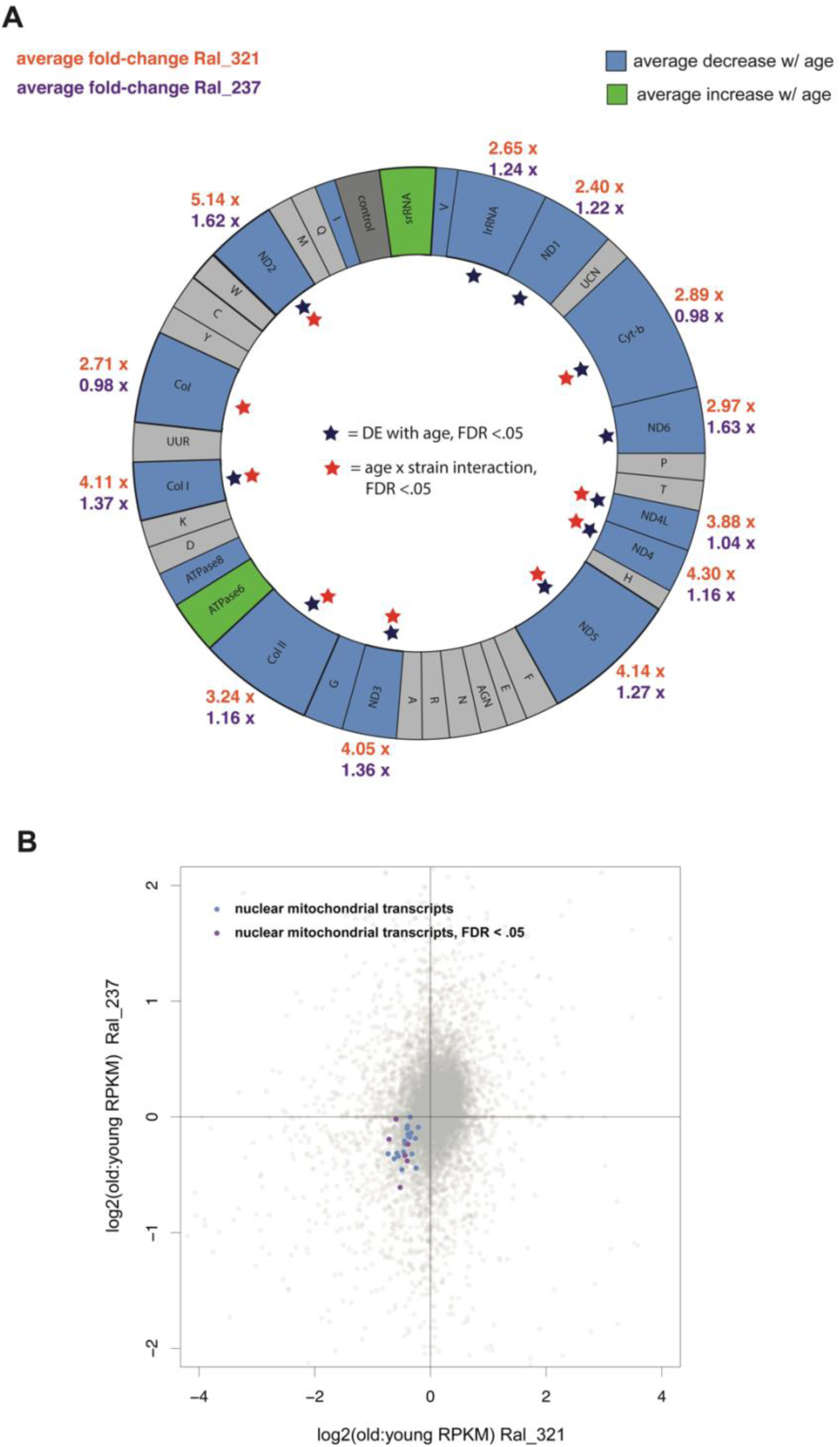
Majority of mitochondrial genome and nuclear mitochondrial transcripts decrease expression in egg chambers with age. (A) There is an average reduction in mitochondrial genome transcript expression in stage 14 egg chambers across strains. Some transcripts are also significant for an age by strain interaction with greater age-related fold-changes (RPKM) in Ral_321. Gray color signifies no expression or no concerted change across strains. (B) Log2 ratios of old to young (RPKM+.5) expression between strains of mitochondrial transcripts from the nuclear genome.

### Downregulation of egg shell chorion transcripts in aged egg chambers show both shared and strain-specific effects

We found a significant gene ontology (GO) enrichment for differentially expressed transcripts associated with eggshell chorion assembly (FDR q-value < 1.44E-04, 15.4-fold enrichment). All of these transcripts were downregulated with age in both strains (Figure 3.9). The downregulation of eggshell transcripts was especially striking in Ral_321, in which all but two eggshell transcripts showed a decrease with age (sign test: p-val < 1.60e-11; Fig 3.3). Ral_237 also showed more eggshell transcripts downregulated with age than expected by chance (sign test p-value <.04) but the effect was not as strong as in Ral_321 (Fig 3.9).

**Figure 3. 3.**
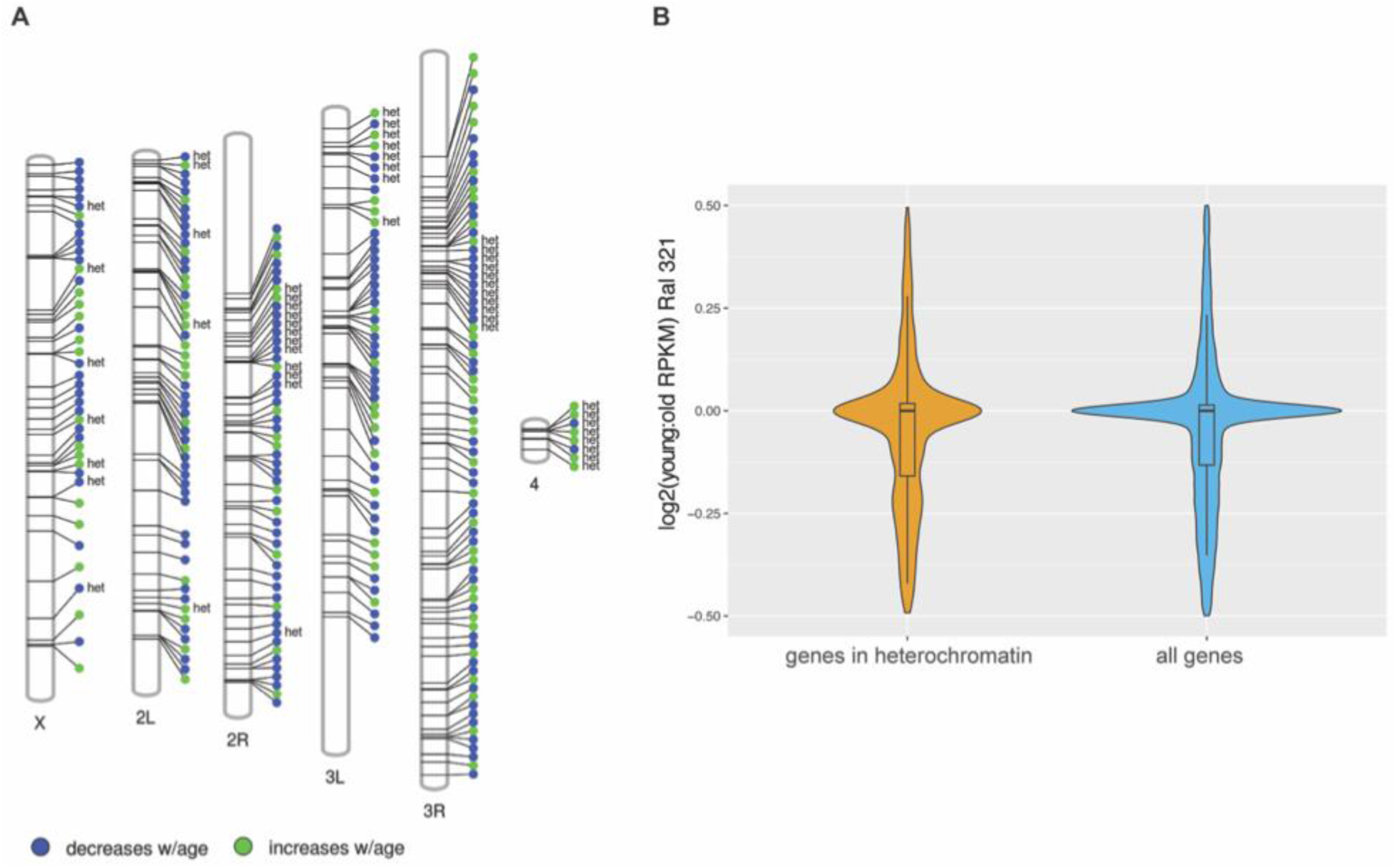
DE transcripts enriched for intercalary heterochromatin and the 4th chromosome. (A) Positional information of differentially expressed genic transcripts across both strains. The notation “het” indicates that the genic location intersects with heterochromatin-associated proteins, H3K9me2/me3, as reported in Kharchenko et al 2011. DE (differentially expressed) genes located in regions of intercalary heterochromatin do not show a concerted directionality change of expression with age (Chi-squared with Yate’s correction, two-tailed p-value <.034). The 4th chromosome is highly enriched for DE genes considering its limited gene composition Chi-squared with Yate’s correction, two-tailed p-value < .0001. (B) Log2(young/old RPKM) of all genes located in heterochromatin versus Log2(young/old RPKM) genome-wide expression change with age. Genes in heterochromatin show similar age-related pattern of expression change as the rest of the genome.

Somatic follicle cells work together to build the protective eggshell in oogenic stages 10-14. This process is dynamic, with transcript amounts changing rapidly between stages (Tootle et al., 2011; Yakoby et al., 2008). Due to the dynamism of expression in late stage oogenesis with regards to eggshell formation, we sought to verify that differential expression of chorion genes was not a consequence of different temporal snapshots in the collected samples. Tootle *et al.* (2011) performed a microarray analysis on 150 genes expressed in a stage-specific manner in the last 24 hours of follicle development, delineated by stages 9- 10a, 10b, 12, and 14. This gene expression dataset included 30 previously known eggshell genes, 19 new candidate chorion genes, and other non-eggshell or chorion genes that showed 4-fold changes in expression at late stages of follicle development. Because this gene expression dataset provides an independent temporal profile of gene expression in late stage oogenesis, we cross-checked our young and old egg chamber expression data against the 49 eggshell-specific transcripts. Critically, gene expression in our samples is strongly correlated with expression in stage-14 egg chambers reported in Tootle 2008 (Pearson’s product-moment correlation = 0.85, p-value < 7.80e-15) but not correlated in stages 9-10, 10b, or 12, confirming that we had captured stage 14 egg-chambers in our analysis (Fig. 3.8).

The decrease in chorion transcripts with age corroborates findings of numerous other studies (Carlson et al., 2015; Doroszuk, 2012; Pletcher, 2002) and here we demonstrated that this age-effect can also vary in effect between strains. The discrepancy between the strains could also be due to the fact that we used chronological age for sampling instead of physiological age. Doroszuk et al., 2012 finds that long-lived flies do not experience a typical decline of reproduction function in the later stages of life which may alternatively explain why we didn’t detect as significant of chorion effects in the strain with slightly longer median lifespan (Doroszuk, 2012; Ivanov et al., 2015).

### Differentially expressed genes in egg chambers enriched for residence in dispersed heterochromatin, but no global genome-wide relaxation of heterochromatic silencing

Previous studies have implicated aberrant gene expression changes with age to changes in the heterochromatin landscape in the soma (Bell et al., 2012; De Cecco, 2013a, b; Jiang, 2013; Larson et al., 2012; Shah et al., 2013; Wood et al., 2010). Genome-wide expression data can be utilized as a proxy for heterochromatic changes by assessing whether genes associated with regions of heterochromatin experience age-related changes in expression. Based on previous studies, we hypothesized that genes located near heterochromatin boundaries, specifically near telomeres and centromeres, may be enriched for differential expression in aging. Kharchenko et al., 2011 described a genome-wide chromatin landscape in Drosophila melanogaster based on 9 prevalent combinatorial patterns of 18 histone modifications (Kharchenko et al., 2011). Pericentromeric heterochromatin domains were characterized by high levels of H3K9me2/me3. We intersected locations of our gene set with the heterochromatin regions described in that study. Of the significantly differentially expressed egg chamber transcripts across both strains in age, we found enrichment for genes in locations of intercalary heterochromatin (Figure 3.3A, 47 differentially expressed genes from 1695 genes in heterochromatin, 300 genes differentially expressed from 14289 total genes; Chi-squared with Yate’s correction, two-tailed p-value = 0.034). We also found a striking enrichment for differentially expressed genes on the fourth or “dot” chromosome, which is primarily heterochromatic and carries only 84 genes (8 genes differentially expressed from 84 total genes on the dot, 300 genes differentially expressed overall from 14289 total genes; Chi-squared with Yate’s correction, two-tailed p-value < .0001). Other than the enrichment for genes on the dot chromosome, there was no obvious signature of enrichment for differentially expressed genes specifically in pericentric heterochromatin (Fig. 3.3A). Critically, we find that the nature of expression change with genes associated with heterochromatin is not in one direction. Differentially expressed genes associated with heterochromatin both increase and decrease during aging (Fig. 3.3A). This is unexpected under the standard heterochromatic aging hypothesis where heterochromatin function becomes lessened and heterochromatic genes become derepressed. Therefore, while heterochromatic regions of the genome tend to be enriched for genes that change in expression during aging, this indicates a general release of regulation, but not release from silencing *per se*.

To test whether there was also a subtle derepression of genes located in heterochromatin genome-wide, we compared age-related expression of all genes which overlapped with heterochromatin in the genome. We found no obvious change in distributions of gene expression ratios between young and old egg chambers of genes located in described regions of heterochromatin compared to the rest of the genome (Figure 3.3B)

We also tested whether the strain specific age-related changes for genes in intercalary heterochromatic regions were due to euchromatic TE insertions that differed between strains. It has been shown that some euchromatic TE insertions can nucleate heterochromatin formation through piRNA targeting (Sentmanat and Elgin, 2012; Shpiz et al., 2014). We used the DGRP strain-specific TE insertion data from TIDAL-fly (Rahman et al., 2015) to compare TE insertion locations across the two strains. However, we did not see strain-specific differences in TE insertions that correlated with aging effects that varied between the two strains.

Other studies have reported decreased expression in heterochromatin modifiers with age. We therefore determined whether genes associated with the gene ontology term for chromatin modifiers showed enrichment for a certain directionality change with age. In egg chambers of both strains, chromatin modifiers tended to increase in expression with age (Ral_237 exact binomial test, p-value < 0.0004; Ral_321 exact binomial test, p-value < .002). Chromatin modifiers in embryos, however, tended to decrease in expression with maternal age (exact binomial test, p-value < 0.03).

### No global release of transposable element expression in aged egg-chambers

Previous studies have shown that transposable elements become derepressed in the soma during aging, notably in brains and fat body of *Drosophila*, and in a variety of other organisms including mammals (Chen et al., 2016; De Cecco, 2013a, b; Li et al., 2013; Maxwell et al., 2011; Patterson et al., 2015). However, a recent study on sequencing artifacts have called some of these results into question (Treiber and Waddell, 2017). Because TEs and small RNA mechanisms of genome defense are primarily expressed in the germline, we aimed to determine whether TE de-repression during aging occurs in reproductive tissues in which they are primarily active. In contrast to other studies, we found no global TE derepression in egg chambers. While one transposable element, *copia*, increased with age across both strains, the other four TEs that showed differential expression with age across strains decreased in expression (Fig. 3.1A, Fig. 3.4, Table 2). Additionally, two TEs, *pogo* and *Juan*, showed a significant strain-by-age effect, exhibiting opposing directions of expression with age across the strains (Fig 3.4C and 3.4D). Figure 3.4D also illustrates that the TEs that are significant in Ral 321 are dispersed throughout the wider distribution of TE expression for Ral 237. There is also no correlation between the ratio of TE expression between young and old egg chambers across strains (Figure 3.4E).

**Figure 3. 4.**
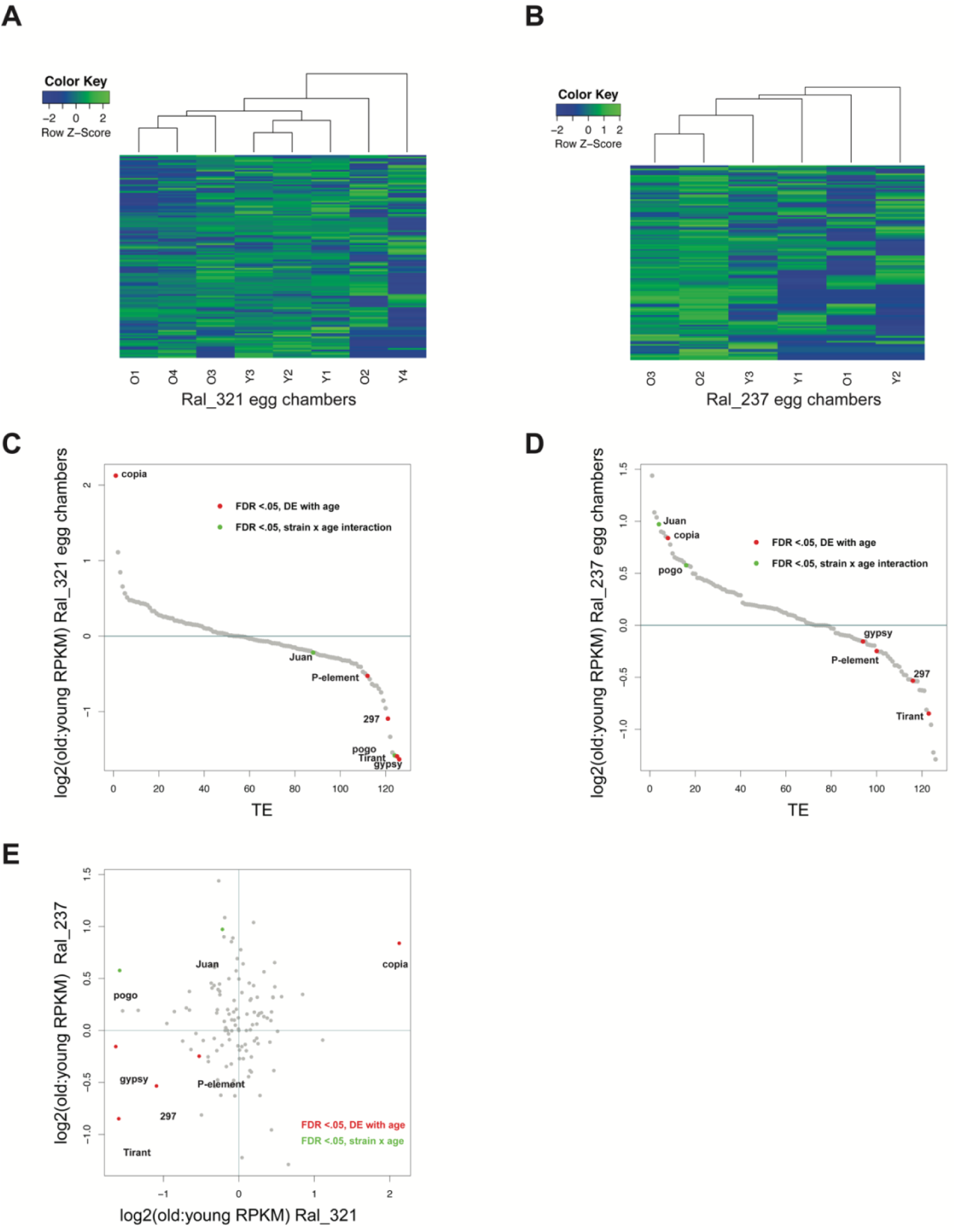
No global derepression of TEs in egg chambers from aged females. A and B) Heatmap of transposable element log2 (RPKM+1) expression in egg chambers normalized by Row Z-Score.“O”= 30-34day old egg chambers, “Y” = 3-4 day old egg chambers. No clear patterns of TE expression occur with age. (C) TEs ordered by ratios of expression from old to young egg chambers in Ral_321. TEs significantly differentially expressed with age in Ral_321 tend to decrease with age. (D) TEs ordered by ratio of expression in Ral_237. Ral_237 shows differentially expressed TEs intercalated through a broader distribution of TE expression. (E) Log2 ratios of old to young RPKM+.5 of TE expression do not show a correlation with age across strains. Two TEs, pogo and Juan show significant age by strain interactions.

**Table 3. 2.** Differential expression results for TEs. TEs that show significant differential expression with age in egg chambers. Two TEs show a strain by age interaction. Fold change refers to fold change differences in RPKM levels. TEs significantly differentially expressed in egg chambers decrease with age in embryos but are not statistically significant.

| *TE* | *EGG CHAMBER* |  |  |  |  | *EMBRYO* |  |
| --- | --- | --- | --- | --- | --- | --- | --- |
|  | fold change |  |  | FDR adj. p-value |  | fold decrease | FDR adj. p-value |
|  | *Ral_321* | *Ral_237* | *drxn w/ age* | *age* | *strain x age* | *Ral_321* | *age* |
| *Tirant* | 3.02 | 1.80 | down | **0.007** | 0.940 | 1.62 | 1.0 |
| *297* | 2.13 | 1.45 | down | **0.010** | 0.869 | 1.78 | 1.0 |
| *Gypsy* | 3.10 | 1.11 | down | **0.011** | 0.125 | 1.06 | 1.0 |
| *Copia* | 4.36 | 1.79 | up | **0.034** | 0.849 | 1.22 | 1.0 |
| *P-element* | 1.44 | 1.19 | down | **0.041** | 0.939 | 1.39 | 1.0 |
| *Pogo* | 2.99 | 1.49 | down, up | 0.125 | **0.027** | 1.90 | 1.0 |
| *Juan* | 1.16 | 1.96 | down, up | 0.597 | **0.039** | 1.37 | 1.0 |

### piRNA pathway transcripts upregulated in aging egg chambers

TE control by piRNA in the germline has been shown to be sensitive to aging. This has been attributed to an increased capacity for TE fragments residing in heterochromatin to contribute to the piRNA pool in older flies (Grentzinger et al., 2012). Moreover, this effect of aging can be transmitted across generations since maternally transmitted piRNA pools establish piRNA biogenesis in offspring. Since some TEs did show significant differential expression with age, we sought to check whether genes in the piRNA pathway, which regulate TE expression in the *Drosophila* germline, showed any age-related-expression changes in egg chambers. Strikingly, 27 out of 31 piRNA pathway genes show an average transcriptional increase with age across the two strains in egg chambers (exact binomial test, p-value < 3.4E-05; Fig. 3.6). piRNA genes are also enriched in the top 10% of differentially expressed transcripts ranked significance (Chi squared with Yate’s correction, p-value = .044). Notably, we did not see these age-effects carried over into the embryo (Fig 3.6C), indicating that this effect may primarily be happening in the follicle cells.

### Differential expression in the aging egg chamber is driven by both somatic and germline changes

Stage-14 egg chambers consist of a mixture of somatic follicle cells and germline material. It is challenging to tease-apart intrinsic aging of the germline from extrinsic factors such as functional decay of the niche (Zhao et al., 2008). We found that *tsunagi* significantly decreased with age in egg chambers (1.75-fold decrease, FDR p-value < 0.0008) (Supp. Table 3.2). *Tsunagi* is required in the germline for proper oogenesis and plays a critical role in in oocyte fate (Mohr et al., 2001; Parma et al., 2007). However, it’s possible that transcript change observed in stage 14 egg-chambers could be coming from the somatic follicular sheath and therefore not necessarily indicative of possible age effects in the oocyte.

We sought to determine whether differential expression during aging was mostly driven by somatic or germline transcripts by performing RNAseq of 0-1 hour embryos of young and old mothers. Maternal germline transcripts can be sequenced because *Drosophila* embryos do not undergo zygotic transcription for approximately two hours. The age-related changes we had seen for *tsunagi* in egg chambers showed the same directionality changes between embryos of young and old mothers (Supp. Table 3.2) supporting the idea that this was an age-related effect occurring in the germline. We next checked whether the Differentially expressed transcripts in egg chambers showed similar directional changes in embryos of Ral_321 as egg chambers across age. It is important to note that many transcripts are not maternally deposited and therefore not expressed in 0-1 hour embryos. The genic transcripts that were differentially expressed in egg chambers and also expressed in 0-1 hr embryos (threshold expression set at above 1.0 RPKM average) did show a significant positive correlation (Pearson’s Product-Moment Correlation, R= 0.31, p-value < 2.85e-06), indicating that some age-related gene expression changes were occurring in the germline rather than simply the follicle cells of stage-14 egg chambers (Fig. 3.7). However, the mitochondrial transcripts with detectable expression in embryos did not show any correlation with changes observed in egg chambers and showed opposite directionality of expression. Thus, we can attribute the observed decrease in mitochondrial transcripts in stage-14 egg chambers to effects in the somatic follicle cells. This also indicates opposing age-related effects occurring in the somatic follicle sheath versus the germline.

We also identified fewer genes with differentially expressed in 0-1 embryos of young versus old mothers when compared to the egg chamber data, though this may be attributed to lower power from fewer replicates. The dynamic nature during early embryogenesis may have also contributed to high variability in gene expression in this stage (Supp. Table 3.2). In 0-1 hour embryos, zygotic transcription is low, but de-adenylation of maternal transcripts may contribute dynamically to rapid changes in apparent gene expression across 0-1 hours.

Our results indicate that age-effects occur in both the somatic cells surround the developing egg as well as in the germline. Additionally, some of transcriptional changes we identify, such as *tsunagi*, may be contributing factors to compromised germ cell division and differentiation that occurs with age.

### Subtle de-repression of TEs in pre-zygotically active embryos of aged mothers

Because TEs are primarily active in the germline and maternal transcripts are deposited into the embryo, we may expect to see a correlation between TE age effects in the egg chambers and embryos. We find no such correlation in expression between late stage egg chambers and 0-1 hour embryos of the same strain (Fig. 3.5B and 3.5C). The TEs that were differentially expressed in egg chambers are for the most part in the middle of the distribution for TE expression in embryos (Figure 3.5B). We do however find a subtle, yet significant enrichment for TEs increasing in expression in embryos of old mothers (Figure 3.5A, Exact binomial test, p < 1.462e-08; Figure 3.5B). The differentially expressed TEs that we observed in egg chambers may be primarily driven by the somatic follicle cells, masking the subtle increase of expression in TEs of the oocyte. Alternatively, there may be an independent de-repression of TEs that occurs in embryos.

**Figure 3. 5.**
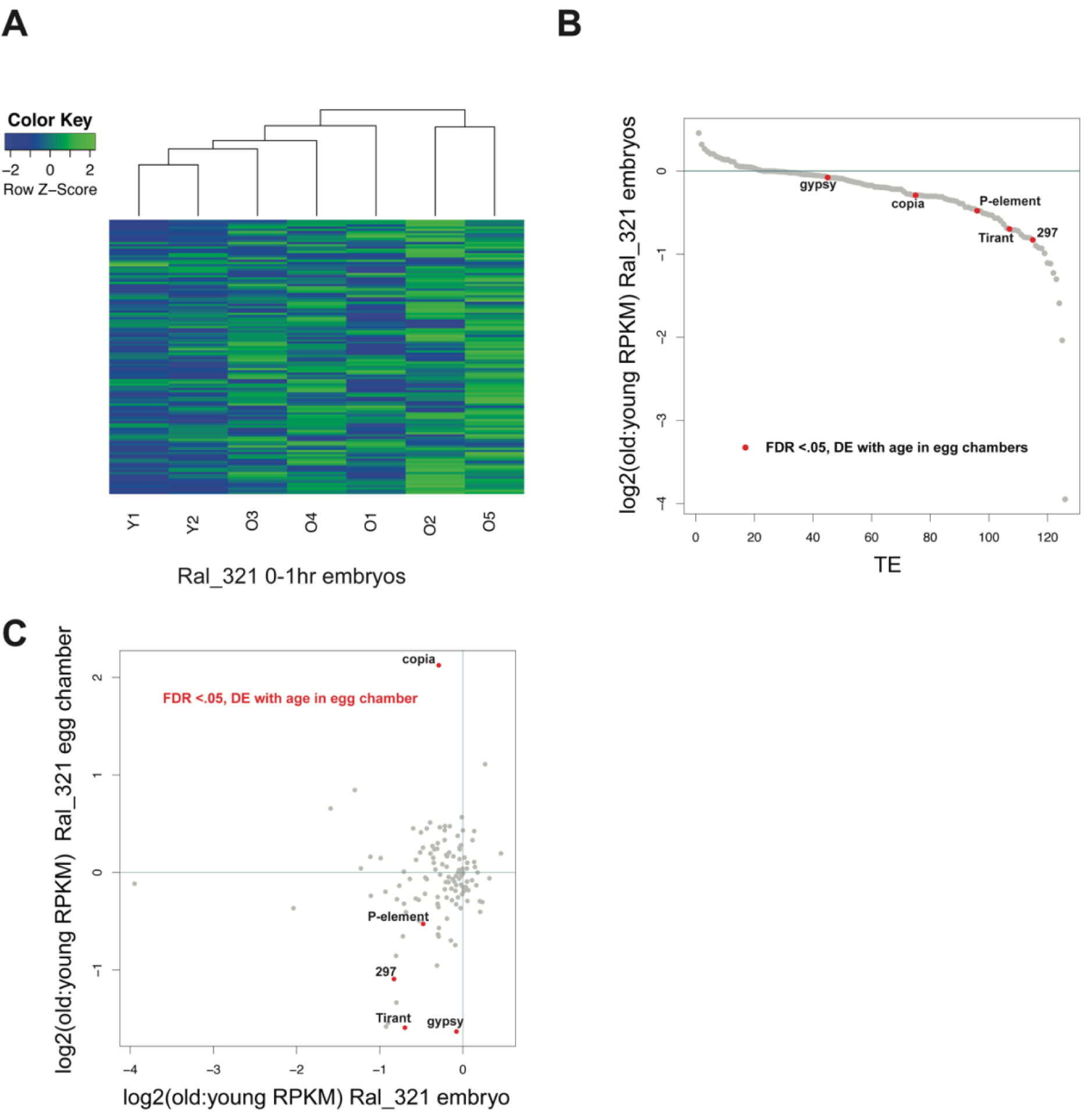
TE age effects differ from egg chamber to embryo. (A) Heatmap of transposable element log2 (RPKM+1) expression in Ral_321 embryos normalized by Row Z-Score. “O”= embryos of 30-34day old mothers, “Y” = embryos of 3-4 day old mothers. Old samples show subtle increase of expression with age. (B) TEs ordered by ratios of expression from embryos of old versus young mothers in Ral_321. Transcripts that were differentially expressed in egg chambers of the same strain are interspersed within the broader distribution of TE expression. The majority of TE transcripts show increased expression with age. (C) Log2 ratios of old to young RPKM+.5 expression between egg chambers and embryos. TE transcripts change in egg chambers are not predictive of TE transcript changes in embryos.

**Figure 3. 6.**
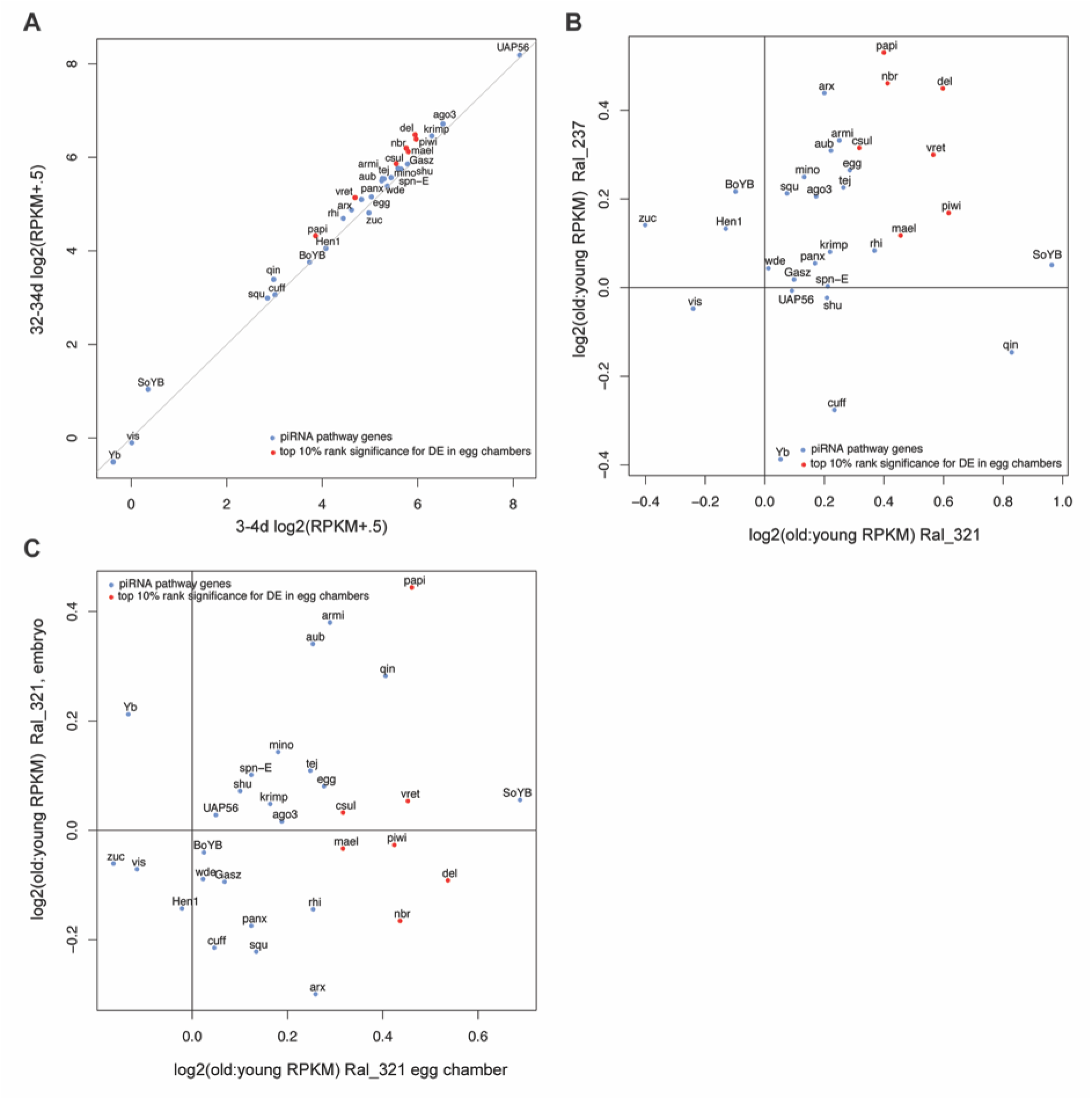
piRNA transcripts increase with age in egg chambers. A) Expression (RPKM+.5) of piRNA pathway transcripts between egg chambers of young and old females. Red dots indicate transcripts that were in the top 10% of significant FDR-adjusted p-values. B) Log2 ratios of old to young piRNA pathway transcript expression (RPKM+.5) in egg chambers across strains. Both strains show a that a majority of piRNA transcripts increase with age. C) Log2 ratios of old to young piRNA pathway expression between egg chambers and embryos. Embryos of old mothers do not show increased transcript expression of piRNA genes.

**Figure 3. 7.**
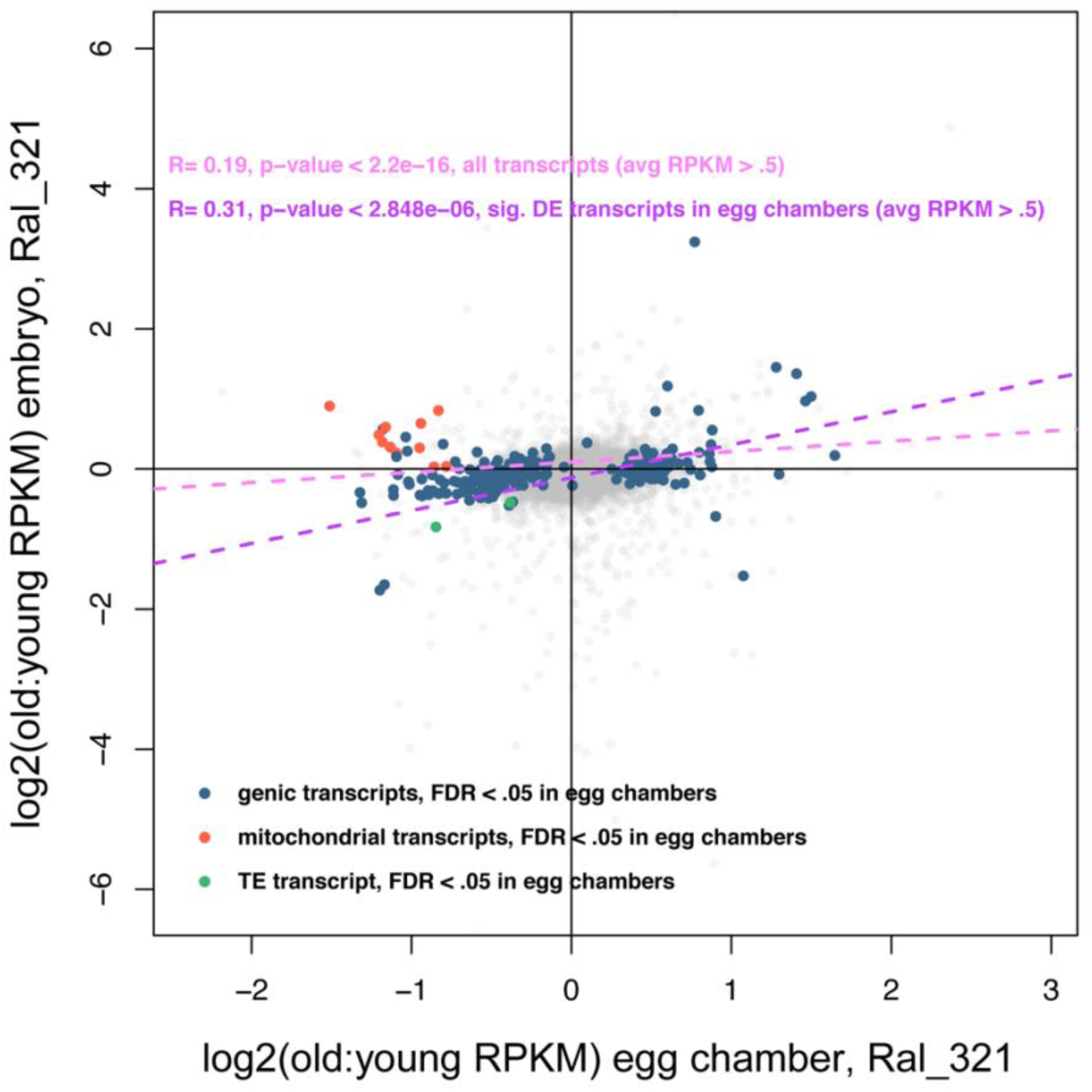
Some age-effects maternally deposited through germline. Log2 ratios of old to young (RPKM+.5) expression between egg chambers and embryos of the same strain. Transcripts that have expression above an RPKM expression threshold of 0.5 in embryos are mildly correlated in age-related change. Transcripts from the mitochondrial genome do not show correlated age-related change between egg chambers and embryos.

**Figure 3. 8.**
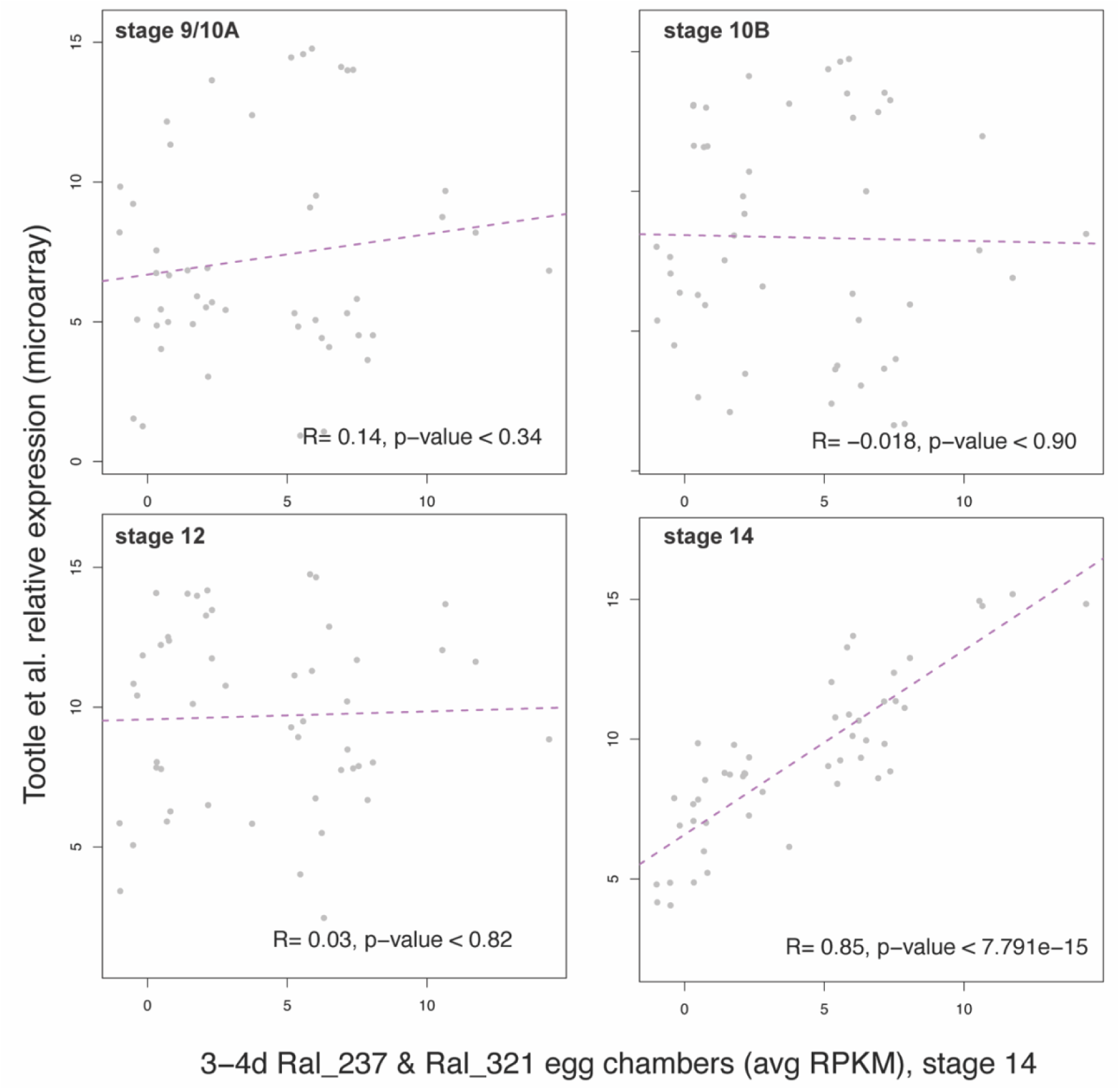
Verification of stage 14 transcript expression. Transcripts that show stage-specific expression in final stages of oogenesis as defined by Tootle et al 2011. Transcript expression from stage 14 egg chambers is strongly correlated with stage 14 oogenic-specific transcript expression but not with the other stages in Tootle et al., 2011.

**Figure 3. 9.**
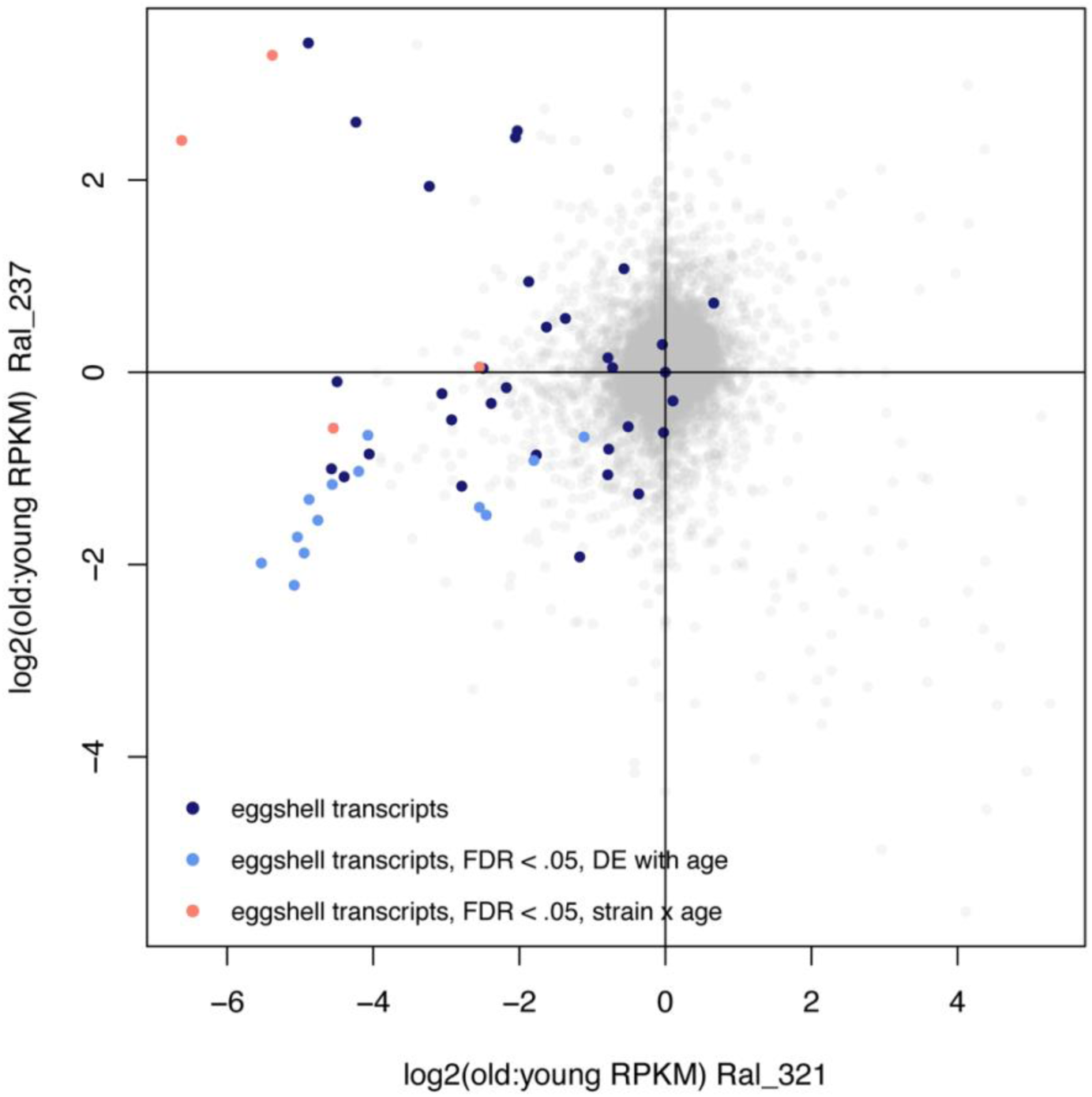
Transcripts associated with the eggshell are downregulated with age in both strains but show stronger age effects in Ral_321. Log2 ratios of expression (RRKM + .5) of transcripts associated with the eggshell between young and old egg chambers across strains.

## DISCUSSION

With delays in childbearing on the rise, the study of reproductive decline grows increasingly relevant (Billari et al., 2007). Fruit flies are an excellent model organism to study because they experience a clear reproductive decline, existing age-related literature in flies is vast, and *Drosophila* share several mechanisms and pathways in ovulation and gametogenesis with mammals (Sun and Spradling, 2013).

Genome-wide RNAseq studies have shown that different tissues vary in age-related signatures, highlighting the importance of analyzing each tissue individually in each species (Sharov et al., 2008; Zhan et al., 2007). Reproductive tissues are unique in that they are a mix of interacting somatic and germline tissue. While the germline is widely recognized as being more resistant to aging than somatic cells, some age-related changes are known to occur. Critically, the relative contribution of factors intrinsic versus extrinsic to the germ line in reproductive decline remains poorly understood.

Here, we report a set of genes that show concerted changes across genetic backgrounds in aged egg-chambers. We additionally highlight the role genetic background plays in age-related effects. For example, while the decline we show in chorion-related transcripts with age parallels other studies, we propose that the severity of this age-effect depends on genetic background.

We also show that aging in late stage egg chambers mirrors that of other tissues, with a downregulation of transcripts from the mitochondria and nuclear transcripts associated with mitochondrial activity. Oocytes have significantly more mitochondria than any other cell, highlighting the incredible energy demands at stake in gametogenesis (May-Panloup et al., 2007). The dysfunction of oocyte mitochondria has been proposed as a possible mechanism involved in reduced competence of oocytes in older human infertility patients (Zhang et al., 2017). One of the most well documented age-effects thought to reduce female fertility is chromosome abnormality in oocytes. There is evidence that reduced mitochondrial activity may contribute to this decline, as improper chromosome segregation has been induced in oocytes deficient in mitochondrial enzymes that metabolize pyruvate (Johnson et al., 2007). Our results support the idea that mitochondrial age effects could contribute to reproductive decline. Because mitochondria are maternally transmitted, the possible deposition of abnormal mitochondria with advanced age has been hypothesized to negatively contribute to offspring health. Here we find no evidence that mitochondrial transcript decline is propagated, as embryos of young and old mothers do not display the same expression patterns as seen in egg chambers. In contrast, maternally deposited mitochondrial transcripts in embryos increase with age. Thus, we propose that the effects of aging in the *Drosophila* ovary on mitochondrial gene expression are largely born out in somatic follicle cells.

Epigenetic changes have been implicated as playing an important role in the aging process in cells of the soma across model organisms. Specifically, genome-wide heterochromatin redistribution during aging has been linked to the de-repression of transposable elements and an overall loss of gene regulation. Whether or not epigenetic factors are perturbed in reproductive and germline tissues is of particular interest because some epigenetic factors are known to transmit across generations (Greer et al., 2016; Grentzinger et al., 2012; Zenk et al., 2017). Current theories of evolutionary aging depend on the assumption that age-related effects do not manifest in offspring. If age-related effects are in fact transgenerational, this could complicate current evolutionary explanations for aging.

While several studies have reported aberrant gene expression in aging on a genome-wide scale (De Cecco, 2013a; Jiang, 2013; Shah et al., 2013), we report no overall loss of gene regulation in aged egg chambers, consistent with another *Drosophila* study using whole bodies (Pletcher, 2002). Previously, it was shown that reporter genes residing in heterochromatin regions of the fly experienced loss of silencing with age (Jiang, 2013). In line with these findings, we too show that genes that show significant age-related expression differences are enriched for regions of heterochromatin, providing evidence for age-related epigenetic changes occurring in late stage egg chambers of *Drosophila* oogenesis. However, we do not however find evidence that the landscape of heterochromatic silencing is relaxed in older egg chambers. Some studies have also reported a decrease in expression of transcripts involved with heterochromatin modification. Here we find that these transcripts do significantly change with age, although in opposite directionality between egg chambers and embryos. This opposing effect in the soma versus germline indicates that patterns of aging may not be universal across tissue types.

Of significant interest is the conserved age-related changes we found between egg chambers and embryos of aged females. These indicate changes in the aged germline *per se*, not simply in the gonad that is a mixture of somatic and germline tissues. These also indicate that aging effects on gene expression in older mothers can be deposited into embryos and transmitted across generations. Since many of the maternal RNA transcripts deposited in embryos are required for embryonic development, this raises the need for further studies of how the maternal transcript pool may change with age and how faithfully those transcripts are deposited into embryos.

A decline in repressive heterochromatin with age has been associated with TEs becoming active and mobile in aging somatic cells (Li et al., 2013; Patterson et al., 2015). Because increased transposition promotes DNA damage and increased mutagenesis, age-related transposable element de-repression has also been proposed to be an important component of genomic instability and a contributor to the prevalence of disease that accompanies advanced age. Here, we find no evidence that TEs are derepressed as a general feature of aging in egg chambers. In contrast, we find that the handful of TEs that are differentially expressed with age tend to decrease in expression with age, in conflict with current TE aging theories, but in line with the idea of adaptive piRNA-mediated immunity with age (Khurana et al., 2011a). The increase in expression in piRNA pathway genes reported here also lends support to this hypothesis and suggests that, in contrast to non-reproductive tissues, mechanisms that limit the harm of TEs may be increased in aging reproductive tissues. It would be worth comparing relative piRNA levels complementary to these TEs in a future study. We also demonstrate that TE age-effects in egg chambers depend on both the genetic background and TE. It is also worth noting that a recent study demonstrates the role artifacts play in leading to incorrect transposition estimation in the soma, possibly throwing previous age-related results into question (Treiber and Waddell, 2017). One interesting finding in our study that deserves further investigation is the subtle increase in TE expression we found when comparing embryos of young and old mothers. In a future study, it would be worth repeating this experiment, paired with a comparison of piRNA profiles of embryos of young and old mothers.

In summary, here we show that there is evidence for age-related change within the reproductive tissues and germline of *Drosophila melanogaster*. However, these tissues are more robust to age-related change in gene expression than the soma, as we find no global TE derepression or global relaxation of heterochromatic silencing with age. We also report that some significant age-related changes in the egg chambers of ovaries persist in embryos. This study supports the conclusion that while there exists a potential to pass on age-related maternal effects, the germline is generally robust to age-related epigenetic changes.

